# Transcriptome-informed brain cartography of polygenic risk and association with brain structure in major psychiatric disorders

**DOI:** 10.1101/2025.05.06.652361

**Authors:** Alessio Giacomel, Timothy R. Powell, Rodrigo R. R. Duarte, Giovanna Nordio, Steve C. R. Williams, Federico Turkheimer, Mattia Veronese, Daniel Martins, Danai Dima

**Author notes:** **Corresponding author:** Alessio Giacomel, Department of Neuroimaging, Centre for Neuroimaging Sciences, Institute of Psychiatry, Psychology and Neuroscience, King’s College London, Denmark Hill, SE5 8AF, London, UK. Equal contribution (Daniel Martins, Danai Dima).

## Abstract

Psychiatric disorders are complex, polygenic conditions characterized by patterned structural brain alterations. Whether these changes reflect transcriptional dysregulation driven by genetic risk remains unclear. We introduce a novel imaging-transcriptomics framework that integrates transcriptome-wide association studies (TWAS) with brain transcriptomic atlases to predict macroscale structural brain abnormalities across seven disorders: ADHD, ASD, AN, BD, MDD, OCD, and schizophrenia (SCZ). We generated disorder-specific Gene Expression-based Disorder Associated Risk (GEDAR) maps and assessed their spatial correlation with observed brain alterations thereby establishing a structured approach to map polygenic transcriptional risk onto macroscale brain phenotypes. We found significant transcriptomic-anatomical correlations in MDD (cortical and subcortical), SCZ (subcortical), and ADHD (subcortical), indicating that regional transcriptional vulnerability might contribute to varying extents to the anatomical expression of genetic risk in these disorders. Pathway enrichment analysis on genetically predicted differentially expressed genes for those disorders where we found spatial correlations between GEDAR maps and observed structural changes revealed immune-related processes as dominant in MDD and SCZ, and neurodevelopmental pathways in ADHD. ASD, AN, OCD, and cortical SCZ lacked significant associations. Importantly, spatial transcriptomic-anatomical alignment did not scale with between-disorder differences in heritability, pointing instead toward additional influences like developmental timing or environmental interactions. These findings underscore the potential and limitations of imaging transcriptomics as a framework for bridging the gap between genetic architecture and systems-level brain changes in psychiatric disorders.

## Introduction

Psychiatric disorders are complex brain conditions with multifactorial aetiologies, arising from the interplay between genetic, environmental, and neurodevelopmental factors^1^. Traditional research approaches have often examined either genetic susceptibility or neuroanatomical alterations in isolation. However, growing evidence suggests that integrating these levels of analysis could yield deeper insights into the biological mechanisms underlying psychiatric conditions^2–5^.

Genome-wide association studies (GWAS) have advanced our understanding of the genetic architecture of mental illness by identifying hundreds of risk loci across disorders. The Psychiatric Genomics Consortium (PGC), established in 2007, has been instrumental in these efforts. Aggregating data from over 400,000 individuals across 36 countries^6^, these studies have unravelled the highly polygenic nature of psychiatric disorders by showing that many common variants of small effect contribute to overall liability. At the same time, neuroimaging techniques have emerged as powerful tools to investigate brain structure and function in mental illness, informing new system-level models of pathology grounded on brain biology. Large-scale efforts such as the ENIGMA (Enhancing NeuroImaging Genetics through Meta-Analysis) consortium have enabled harmonized analyses of magnetic resonance imaging (MRI) data across thousands of participants, revealing reproducible patterns of structural brain alterations in various psychiatric conditions^7,8^. These patterns include both disorder-specific and transdiagnostic differences, which are not randomly distributed across the brain but instead suggest regional variation in vulnerability—potentially shaped by intrinsic biological properties^9,10^.

Neuroimaging phenotypes have been increasingly conceptualized as intermediate markers on the path from genes to behaviour^11^. Imaging genetics, a discipline that integrates genomic and neuroimaging data, has uncovered biological pathways linking genetic risk to brain phenotypes^12^. A central assumption of this field is that dysregulated molecular processes—e.g., altered gene expression or protein function—translate into macroscale changes in brain structure and function. These changes, in turn, underlie the behavioural and clinical features of psychiatric disorders^13^. Yet, whether the patterned imaging phenotypes documented in patient groups can be predicted directly from the spatial distribution of genetically-encoded, potentially dysregulated, risk gene expression remains an open empirical question.

A potentially informative way to address this question would be to ask whether regional brain changes observed in large-scale neuroimaging studies, such as those from ENIGMA, can be spatially predicted using knowledge about the **genetically influenced gene expression** and its **constitutive architecture of the human brain**. The availability of the Allen Human Brain Atlas (AHBA)^14^—the most comprehensive resource of spatial gene expression in the human brain—**now offers this possibility**^2^. Imaging transcriptomics is an emergent line of research which investigates whether patterns of brain changes align spatially with regional gene expression, thereby linking molecular and systems-level pathology. Exploratory studies have shown that genes whose spatial expression correlates with brain alterations in disorders such as schizophrenia^3,15^, major depressive disorder^16^, and autism^15^ are often enriched for known disorder-related genetic risk, suggesting that regional brain vulnerability may be rooted in the local molecular landscape^17^. However, the field is still missing a **structured framework to integrate GWAS summary statistics, transcriptomic prediction, and neuroimaging phenotypes** to probe the genetic risk–mRNA transcription–brain anatomy–disorder axis continuum. Studying several disorders in parallel offers a unique opportunity to achieve the later goal given the wide variation in genetic and environmental contributions across conditions. For example, schizophrenia and ADHD exhibit high heritability (∼75–80%), while disorders such as major depressive disorder (MDD) show more moderate estimates (∼35– 45%), implying a larger role for environmental factors^18^. This variability implies that that the degree to which brain changes can be predicted from genetically regulated gene expression dysregulation could scale with a disorder’s heritability, but this remains largely untested.

**In this study**, we sought to address this gap by introducing and testing a cross-disciplinary framework combining bioinformatics, statistical genetics, and neuroimaging methods to investigate whether regional patterns of brain structural change can be predicted from polygenic risk-informed gene expression signatures. We focused on seven main psychiatric conditions with available large-scale genetics and neuroimaging data, including attention-deficit/hyperactivity disorder (ADHD), autism spectrum disorder (ASD), anorexia nervosa (AN), bipolar disorder (BD), major depressive disorder (MDD), obsessive-compulsive disorder (OCD), and schizophrenia (SCZ). First, we developed spatial maps of genetically-predicted gene expression profiles—**Gene Expression-based Disorder Associated Risk (GEDAR)**—by integrating GWAS summary statistics from the PGC with transcriptomic prediction models constructed using S-MultiXcan^19^, to infer disorder-specific regional gene expression profiles across the brain. Next, GEDAR maps were independently spatially correlated with regional cortical thickness and subcortical volume differences observed in ENIGMA case-control meta-analyses. Finally, we examined whether the strength of this spatial correspondence varied systematically with disorder-specific heritability estimates from twin studies and pathway enrichment analysis.

## Methods

### Psychiatric GWAS Datasets

We obtained publicly available GWAS summary statistics from the Psychiatric Genomics Consortium (PGC) for seven psychiatric disorders: attention-deficit/hyperactivity disorder (ADHD), autism spectrum disorder (ASD), anorexia nervosa (AN), bipolar disorder (BD), major depressive disorder (MDD), obsessive-compulsive disorder (OCD), and schizophrenia (SCZ). Details on sample size, ancestry composition, and imputation procedures are available in the corresponding PGC publications listed in Table 1.

**Table 1.**
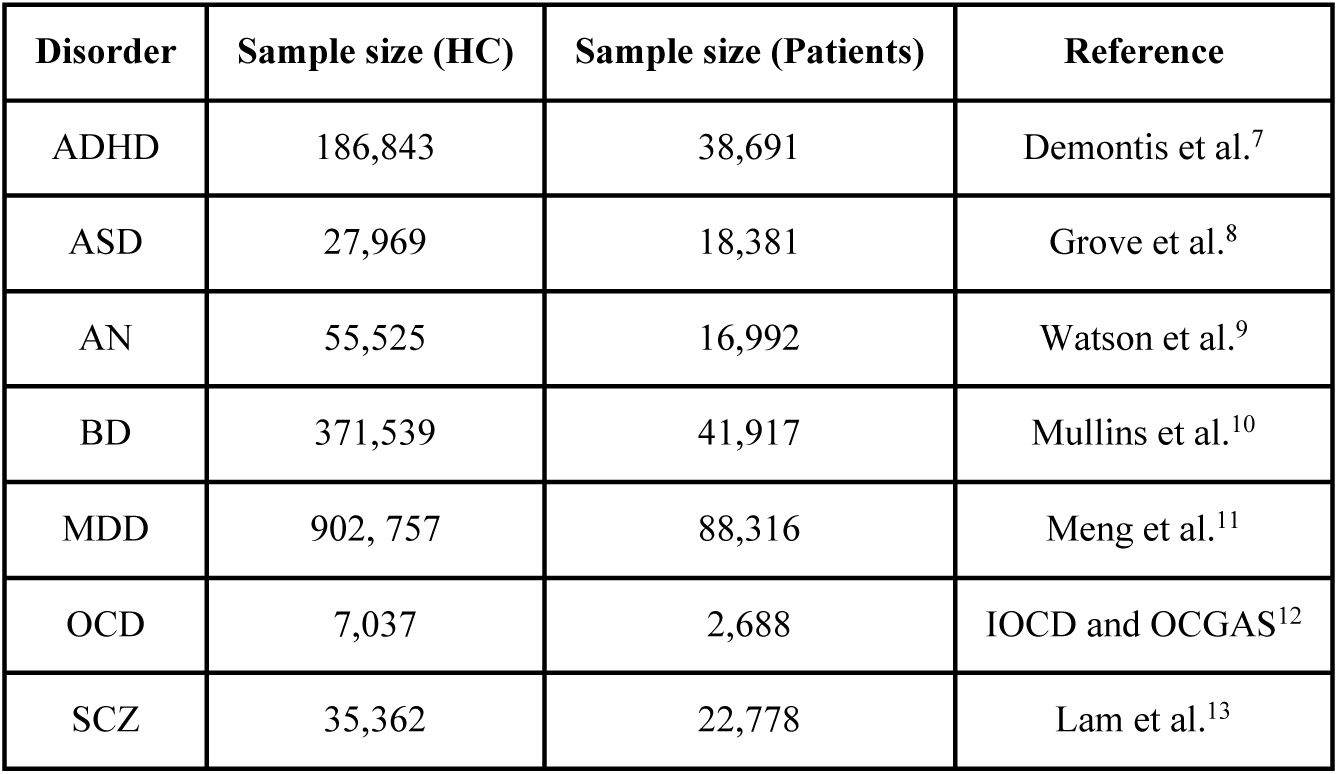
Summary of the sample sizes in the Genome Wide Association Studies (GWAS) from the Psychiatric Genomic Consortium (PGC) used in our analysis (sample size and publication reference).

| Disorder | Sample size (HC) | Sample size (Patients) | Reference |
| --- | --- | --- | --- |
| ADHD | 186,843 | 38,691 | Demontis et al.<sup>7</sup> |
| ASD | 27,969 | 18,381 | Grove et al.<sup>8</sup> |
| AN | 55,525 | 16,992 | Watson et al.<sup>9</sup> |
| BD | 371,539 | 41,917 | Mullins et al.<sup>10</sup> |
| MDD | 902,757 | 88,316 | Meng et al.<sup>11</sup> |
| OCD | 7,037 | 2,688 | IOCD and OCGAS<sup>12</sup> |
| SCZ | 35,362 | 22,778 | Lam et al.<sup>13</sup> |

### TWAS and Gene Expression Prediction

At first, we developed a spatial model of brain structural variation by mapping polygenic risk to genes using H-MAGMA^20^, integrating gene expression data, and correlating dysregulation scores with psychiatric disorder-related cortical and subcortical abnormalities from ENIGMA (for more information see Supplementary Material). This resulted in a list of prioritised genes based on predicted cumulative risk of dysregulation, with no information of change whether genes are down-regulated or up-regulated. Direction of regulation might be an important factor in explaining regional variation in brain structural changes, since upregulated genes could be protective and down-regulated pathological (or vice-versa) leading potential downstream effects on brain structure to be cancelled out. Since direction was a feature we wanted to account for, we turned to Transcriptome-Wide Association Studies (TWAS)^21^ performed using the S-MultiXcan framework to estimate genetically predicted gene expression levels from GWAS summary statistics^19^. S-MultiXcan models trained on the 11 brain tissue panels from the Genotype-Tissue Expression (GTEx) dataset v8^22^ were used to generate gene-level Z-scores representing up- or down-regulation across tissues. S-MultiXcan takes S-PrediXcan Z-scores (or betas) for each gene across multiple tissues and applies a multivariate regression that accounts for cross-tissue correlation, resulting in a single overall Z-score or p-value that reflects the joint association signal across tissues. This was the ideal approach for our study, due to incomplete correspondence between GTEx tissues and regions of the parcellations used in the study (i.e., Desikan-Killiany for cortical regions and aseg for subcortical regions). This strategy also ensured uniform application of gene expression predictions across the entire cortical and subcortical atlas.

### Gene expression and GEDAR Calculation

Regional microarray expression data were obtained from 6 post-mortem brains (17% female, ages 24.0-57.0, 42.50±13.38) provided by the Allen Human Brain Atlas (AHBA)^14^. Regional data were processed with the *abagen* toolbox (version 0.1.3; https://github.com/rmarkello/abagen)^23^ using the Desikan-Killiany (DK) atlas for cortical regions and the aseg atlas for subcortical regions^24,25^. The obtained regional expression matrix comprised 83 rows, corresponding to brain regions (cortical and subcortical), and 15,633 columns, corresponding to the retained genes after filtering out genes with below background expression signal. To increase specificity to genes expressed in the brain, the resulting genes were filtered by the genes identified in the Human Protein Atlas (HPA)^26^ as expressed (i.e., genes that have an expression above cut-off of 1 nTPM).

The Gene Expression-based Disorder Associated Risk (GEDAR) score for region j was calculated using the following formula:

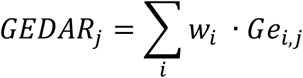

Where *w*_*i*_ indicates the weight for gene *i* as expressed by the predicted TWAS Z-score, and *Ge*_*i*,*j*_ the gene expression for region *j* and gene *i* as expressed by the AHBA. We first analysed the effect of genetic-risk informed TWAS associated signals by using the sign of the Z-score as proxy for the gene directionality (i.e., adding the weighted effect of positive Z-scores and removing the effects of negative Z-scores). Furthermore, to evaluate the effects of gene directionality, we applied two additional weighting schemes where we consider separately *up-* and *down-regulated* TWAS association signals. In the up-regulated scheme, we included only genes with positive Z-scores in the TWAS, while the down-regulated scheme included genes with negative Z-scores only. We then calculated the GEDAR brain maps for all disorders and weighting schemes using three arbitrary thresholds (top 1%, 5%, and 10%) of the TWAS-predicted differentially regulated genes, ranked according to p-value. Lower thresholds also commonly used in the calculation of polygenic risk scores (e.g., 0.1% and 0.5%)^27^ were not used in this study as we noticed that they would result mostly in the selection of only a few single genes, which would be equivalent to a single candidate and not a polygenic gene approach. GEDAR scores were calculated independently for each region of the DK atlas and aseg atlas for subcortical regions to match the regions of the ENIGMA parcellation, thus obtaining a cortical and a subcortical GEDAR brain map for each investigated disorder.

### ENIGMA Structural Brain Changes Reference Maps

We used disorder-specific structural brain maps from the ENIGMA consortium, based on large-scale case–control meta-analyses of MRI data. Briefly, cortical thickness and subcortical volumes (caudate, putamen, pallidum, nucleus accumbens, thalamus, hippocampus, amygdala and lateral ventricles) were extracted using FreeSurfer (http://surfer.nmr.mgh.harvard.edu) from high-resolution T1-weighted MRI brain scans. FreeSurfer segments and parcellates the cortical surface to provide precise measurements across predefined regions of interest (ROIs) based on the DK atlas. For each disorder, we extracted regional meta-analytical Cohen’s *d* effect sizes for cortical thickness and subcortical volumes. These ENIGMA-derived maps served as the neuroimaging reference phenotype for testing GEDAR–brain alignment. Although both cortical thickness and surface area are standardly inspected as outcome metrics by ENIGMA studies, we decided to focus on thickness only as this metric has been shown to present higher regional heritability (thickness: 0.20 – 0.76 vs surface area: 0.03 – 0.75), which means its variation is under stronger genetic control. This would therefore offer larger statistical power to test our hypotheses while keeping the need for multiple testing correction at the necessary minimum^28^. Details on the samples used in each ENIGMA study and respective sources are summarized in Table 2.

**Table 2.** Summary of ENIGMA’s cortical thickness and subcortical volume data used for the study. We provide summary data on the sample sizes, mean age and proportion of female participants, discriminating between groups where data was available. Abbreviations: Attention Deficit and Hyperactivity Disorder - ADHD, Autism Spectrum Disorder - ASD, Anorexia Nervosa - AN, Bipolar Disorder - BD, Major Depressive Disorder - MDD, Obsessive-Compulsive Disorder - OCD and Schizophrenia - SCZ.

| Feature | Disorder | Sample size (HC) | Sample size (Patients) | Age (yrs) | Sex (F) | Reference |
| --- | --- | --- | --- | --- | --- | --- |
| Thickness | ADHD | 1934 | 2246 | 19.22 ± 11.31 | 25.9% | Hoogman et al. 2019<sup>29</sup> |
|  | ASD | 1651 | 1571 | 15.62 ± 8.52 | 19% | Van Rooij et al. 2019<sup>30</sup> |
|  | AN | 963 | 685 | 21 | 100% | Walton et al. 2022<sup>31</sup> |
|  | BD | 4056 | 2447 | - | - | Hibar et al. 2018<sup>32</sup> |
|  | MDD | 7957 | 2148 | - | - | Schmaal et al. 2017<sup>33</sup> |
|  | OCD | 1436 | 1498 | - | - | Boedhoe et al. 2018<sup>34</sup> |
|  | SCZ | 5098 | 4474 | HC: 32.8<br>Patients: 32.3 | HC: 47%<br>Patients: 44% | Van Erp et al. 2018<sup>35</sup> |
| Subcortical volumes | ADHD | 1529 | 1713 | 18.6 ± 11.81 | 34% | Hoogman et al 2017<sup>36</sup> |
|  | ASD | 1651 | 1571 | 15.62 ± 8.52 | 19% | Van Rooij et al. 2019<sup>30</sup> |
|  | AN | 963 | 685 | 21 | 100% | Walton et al. 2022<sup>31</sup> |
|  | BD | 2594 | 1710 | - | 56% | Hibar et al. 2016<sup>37</sup> |
|  | MDD | 7199 | 1728 | - | - | Schmaal et al. 2016<sup>38</sup> |
|  | OCD | 1472 | 1495 | - | - | Boedhoe et al. 2016<sup>39</sup> |
|  | SCZ | 2540 | 2028 | 31 HC<br>34 Patients | 48% HC<br>31% Patients | Van Erp et al. 2016<sup>40</sup> |

### Spatial Correlation Analysis

We calculated Spearman rank correlations between regional GEDAR scores and ENIGMA Cohen’s *d* values for each disorder, separating between cortical and subcortical regions. To account for spatial autocorrelation in cortical data, we applied a spherical spin permutation method (n = 1000 rotations) using the *neuromaps* toolbox (https://github.com/netneurolab/neuromaps)^41^ - Vasa method^42^. Subcortical correlations were tested using standard permutation due to lack of surface geometry. Results were considered significant at *p_spin_* < 0.05.

**Figure 1.**
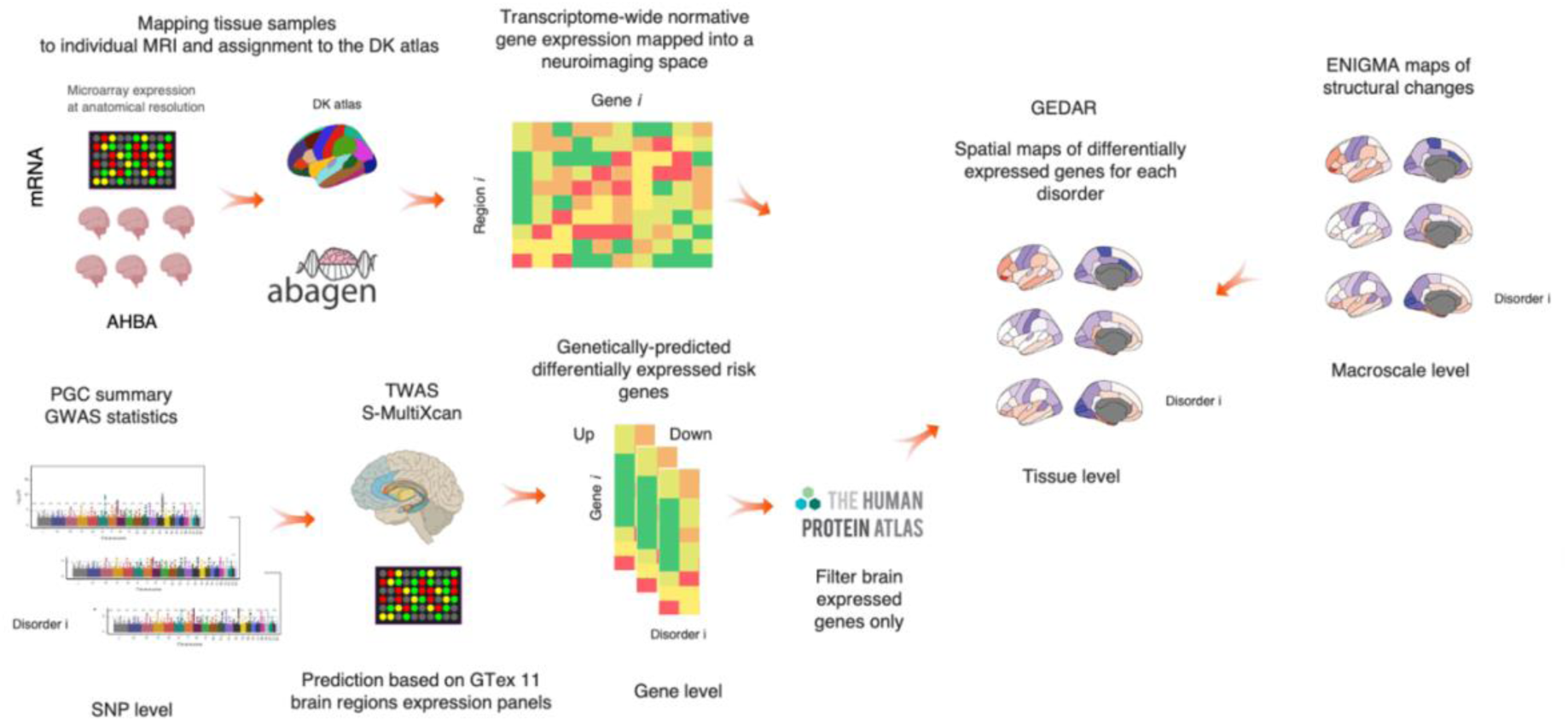
Predicted TWAS differentially regulated genes. Volcano plots depicting differentially regulated genes as predicted by the application of S-MultiXcan to PGC available GWAS summary data from each disorder. The x axis displays TWAS predicted effect sizes (z-scores) and the y axis −log10 (p-values). Colour bar highlights variation in −log10(p-values) within the range of the top 10% most significant genes. Labelled genes represent the top 1%. Abbreviations: Attention Deficit and Hyperactivity Disorder (ADHD), Anorexia Nervosa (AN), Autism Spectrum Disorder (ASD), bipolar disorder (BD), Major Depressive Disorders (MDD), Obsessive-Compulsive Disorder (OCD) and Schizophrenia (SCZ).

### Meta-analytical estimates of differentially expressed genes in post-mortem brain samples

We performed an additional analysis comparing GEDAR brain maps calculated from differentially expressed genes in post-mortem brain samples reported in the literature by Sadeghi et al.^43^. We used the same procedure as previously described for projecting the genes on all regions using the reported effect sizes as weights. The analysis was performed to assess whether genes predicted from TWAS on brain specific tissue models would yield different results from differentially expressed genes identified experimentally from a well-powered meta-analysis of transcriptomic studies in post-mortem brain samples. This additional analysis could not be performed on all disorders investigated here but only on a subset (ASD, BD, MDD and SCZ) as data for the other disorders were not included in the original meta-analytical work from Sadeghi et al.^5^. For the correlation analysis, we used the same procedure described above.

### Heritability in twin-studies

We compiled data on heritability estimates from twin studies for the seven psychiatric disorders included in this study to contextualize the extent of genetic influence across conditions. A summary of these data and respective sources are presented in Table 3. We decided to rely on heritability estimates from twin studies instead of SNP-estimated heritability to avoid circularity as we used GWAS summary statistics to run our TWAS analyses. These differences in heritability provide a useful framework for interpreting the observed variation in the correspondence between genetically predicted transcriptomic risk (GEDAR) and regional brain structural alterations. Specifically, we explored whether disorders with higher heritability show stronger spatial alignment between gene expression-based risk maps and neuroimaging phenotypes. To do so, we computed Spearman correlation coefficients between heritability estimates and GEDAR–ENIGMA correlation values across the seven psychiatric disorders. For each disorder, we used the midpoint of the reported heritability range from twin studies as a summary measure. This analysis was performed separately for each combination of gene regulation scheme (negative, positive, both), inclusion threshold (top 1%, 5%, and 10% of genes), and brain compartment (cortical or subcortical). Spearman correlations and associated *p*-values were computed using the *`scipy.stats.spearmanr`* function in Python.

**Table 3.** Twin-based heritability estimates for seven psychiatric disorders. Heritability estimates (h²) represent the proportion of variance in liability explained by genetic factors, as determined by twin studies. Estimates are presented alongside references to the original source studies. Abbreviations: Attention Deficit and Hyperactivity Disorder - ADHD, Autism Spectrum Disorder - ASD, Anorexia Nervosa - AN, Bipolar Disorder - BD, Major Depressive Disorder - MDD, Obsessive-Compulsive Disorder - OCD and Schizophrenia - SCZ.

| Disorder | Twin-based Heritability Estimate ($h^2$) | Reference |
| --- | --- | --- |
| ASD | 70-90% | Tick et al., 2016<sup>44</sup> |
| SCZ | 75-80% | Hilker et al., 2018<sup>45</sup> |
| ADHD | 70-80% | Faraone & Larsson, 2019<sup>46</sup> |
| BD | 60-85% | McGuffin et al., 2003<sup>47</sup> |
| AN | 50-60% | Yilmaz et al., 2015<sup>48</sup> |
| OCD | 45-60% | van Grootheest et al.,<br>2005<sup>49</sup> |
| MDD | 35-45% | Sullivan et al., 2000<sup>50</sup> |

### Pathway Enrichment Analysis

To explore the biological significance of genes included in each GEDAR map estimation (i.e., TWAS identified genes) for which we found significant association with observed structural changes (by disorder, direction, and threshold), we conducted gene ontology (GO) and pathway enrichment analysis using the ToppGene Suite (https://toppgene.cchmc.org/). We focused on Gene Ontology (GO) Biological Processes (GO:BP), Molecular Function (GO:MF) and Cellular Component (GO:CC). Functional enrichment analysis was performed using the g:Profiler g:GOSt online tool (version e112_eg59_p19_87fff33f, accessed on January 31, 2025), an online tool for functional profiling of gene lists^20^. To account for multiple testing, we applied the g:SCS (Set Counts and Sizes) threshold as implemented by the g:GOSt tool (Reimand et al., 2007). Pathways with a false discovery rate (FDR)–corrected *p* < 0.05 were considered as significantly enriched.

### Data Availability

All GWAS summary statistics used in this study were obtained from the publicly available resources of the Psychiatric Genomics Consortium (https://www.med.unc.edu/pgc). Neuroimaging data were derived from meta-analytic results published by the ENIGMA consortium (http://enigma.ini.usc.edu). Gene expression data were sourced from the Allen Human Brain Atlas (https://human.brain-map.org/). GTEx panels used for the TWAS analyses and methods originating this dataset are available at https://www.gtexportal.org/home/. Meta-analytical estimates of gene differential regulation in post-mortem brain samples and heritability estimates data are available from the corresponding sources cited within the main text and supplementary materials.

### Code Availability

All scripts used for GEDAR computation, spatial correlation analysis, and figure generation are available upon reasonable request to the corresponding author. A cleaned version of the analysis pipeline will be made available on GitHub upon publication to promote transparency and reproducibility.

### Ethics Statement

This study used exclusively secondary analyses of publicly available anonymized data and did not involve the collection of new human participant data. As such, ethical approval and informed consent were handled by the original studies that generated the datasets. All procedures in the original studies were conducted in accordance with institutional and international research ethics guidelines.

## Results

### TWAS Prediction

The TWAS analysis conducted using S-MultiXcan across the 11 GTex brain tissue panels, identified a similar number of differentially expressed genes per disorder, ranging from 2,759 to 2,768: ADHD (N=2,759), ASD (N=2,767), AN (N=2,763), BD (N=2,767), MDD (N=2,764), OCD (N=2,764), and SCZ (N=2,768). Volcano plots depicting expression change in either direction are shown for each disorder in Figure 2. All disorders displayed a characteristic U-shaped pattern, reflecting balanced distributions of up-regulated (positive Z-score) and down-regulated (negative Z-score) genes. Although the overall distribution of gene expression changes was broadly similar across disorders, some showed more pronounced clustering. In particular, AN exhibited SPINK8 as the most significantly down-regulated gene (Z = –5.24, p < 0.001), while OCD was characterized by BRD2 down-regulation (Z = –0.50, p < 0.001). MDD showed NEGR1 as the most significantly up-regulated gene (Z = 5.60, p < 0.001). For SCZ, two distinct gene clusters were observed: an up-regulated cluster (NOTCH4, Z = 5.00, p < 0.001), and a larger down-regulated cluster involving BRD2 (Z = –4.13), AGER (Z = –5.63), HLA-DMB (Z = –7.66), HLA-DMA (Z = –7.96), and CCHCR1 (Z = –6.85), all p < 0.001.

**Figure 1.**
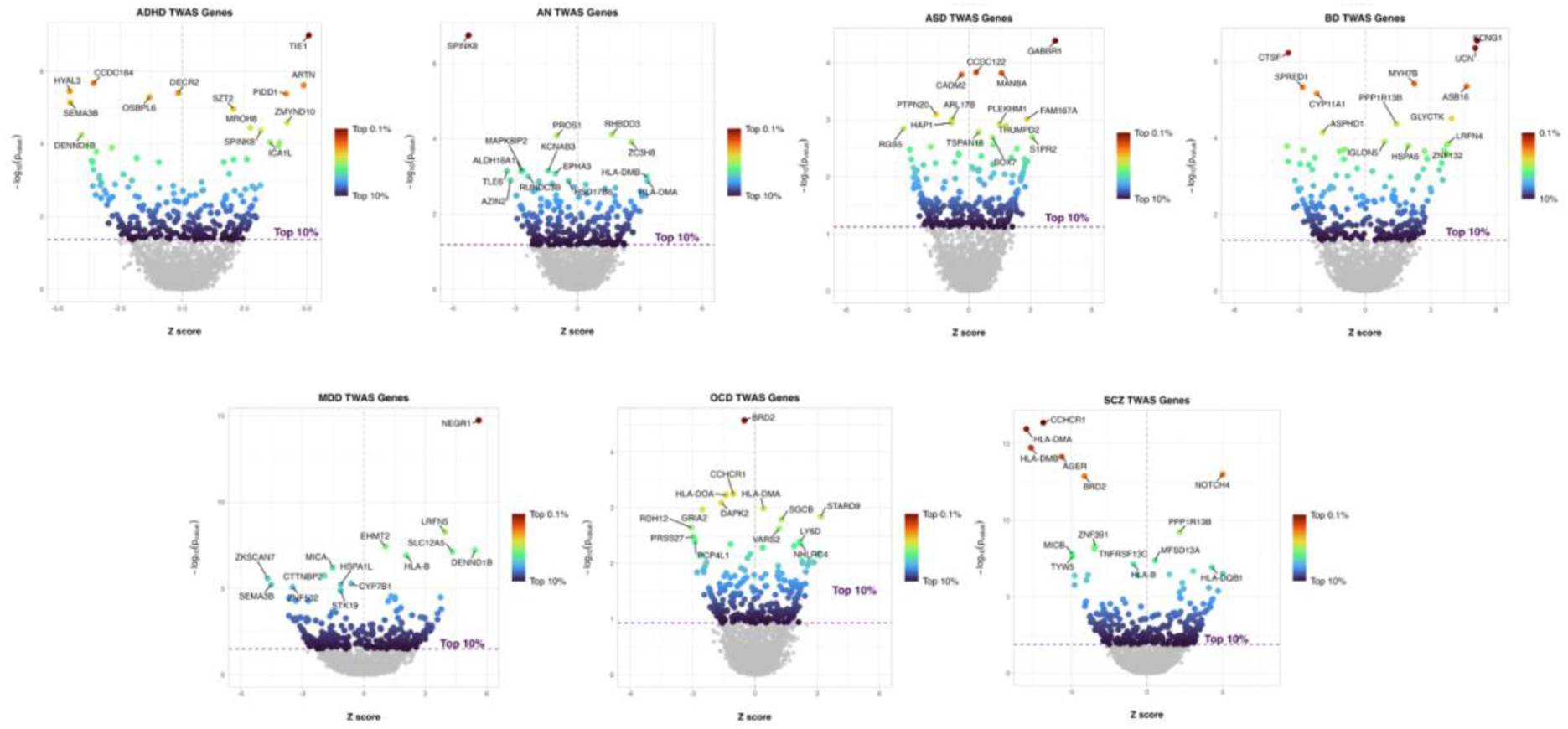
Step-by-step multimodal integration workflow combining genetic, transcriptomic, and neuroimaging data. First, transcriptomic data from the Allen Human Brain Atlas (AHBA) are mapped onto cortical regions defined by the Desikan-Killiany atlas using the *abagen* toolbox, generating region-specific gene expression profiles. Next, transcriptome-wide association studies (TWAS) are performed using the *S-MultiXcan* method, leveraging genome-wide association study (GWAS) summary statistics from the Psychiatric Genomics Consortium (PGC) and predictive models trained on expression data from 11 brain regions in the Genotype-Tissue expression (GTEx) database. Genes are filtered to retain only those expressed in brain tissue, according to the Human Protein Atlas. The resulting genetically predicted gene-expression differences are spatially mapped onto cortical regions using the Gene Expression-based Disorder-Associated Risk (GEDAR) approach, generating disorder-specific cortical expression maps. Finally, these GEDAR-derived expression maps are compared with structural neuroimaging data from the ENIGMA consortium, linking molecular genetic risk factors to observed macroscale structural brain alterations in psychiatric disorders.

**Figure 2.**
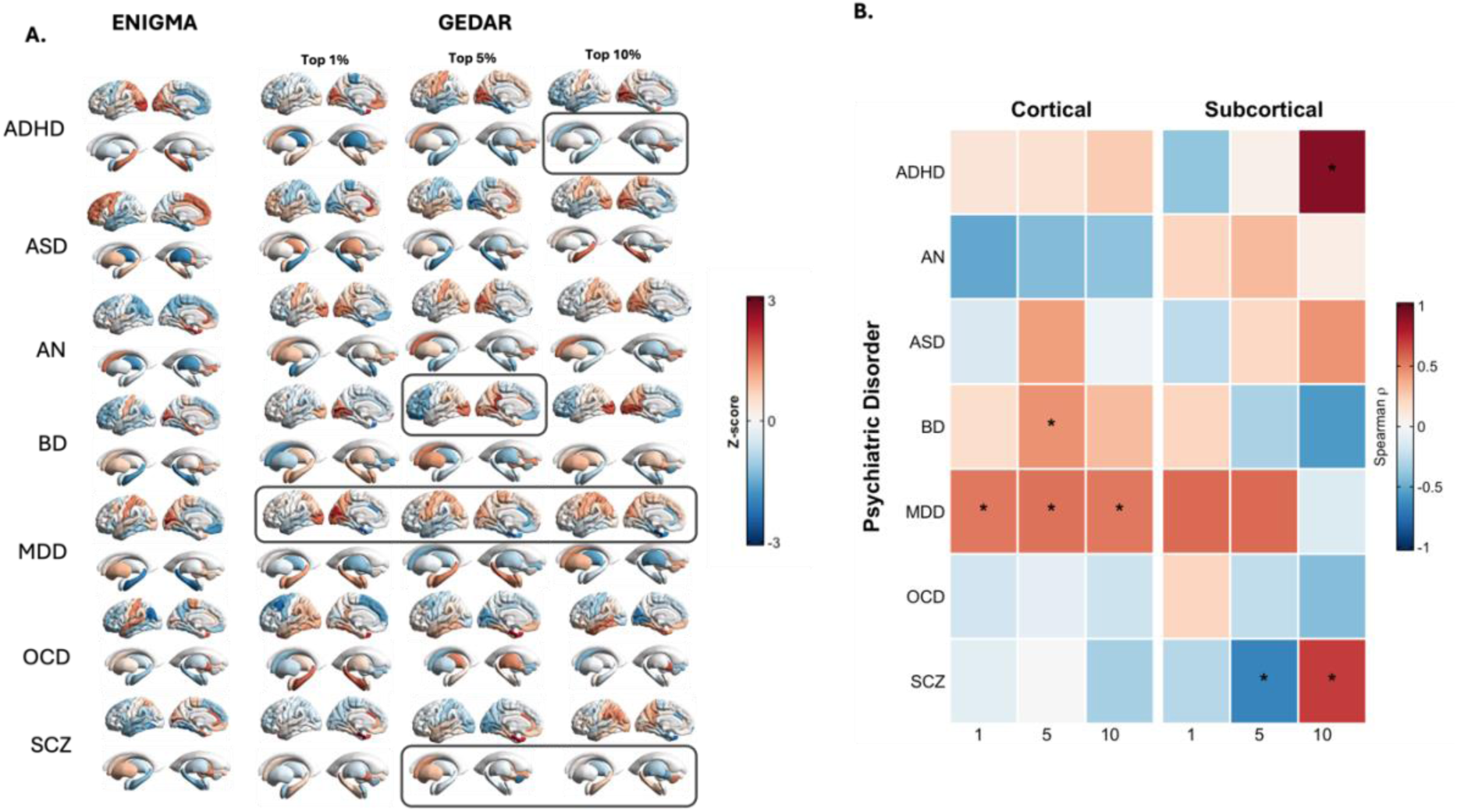
Correlation between ENIGMA and GEDAR brain maps (calculated from both up- and down-regulated TWAS genes). Panel A shows regional distributions of Z-scores of Cohen’s d effect sizes capturing changes in cortical thickness and subcortical volumes between patients and healthy-controls as provided by the respective ENIGMA meta-analysis (left panel); we also display our estimated GEDAR maps estimated from both up- and down-regulated TWAS genes by applying three different thresholds (top 10, 5 and 1%) to select the most differentially expressed genes. Panel B depicts spearman correlations between ENIGMA maps of structural differences and GEDAR maps calculated at the three different thresholds separately for cortical and subcortical regions. Significance was calculated by generating 1000 random spin permutations to account for spatial autocorrelation. The * highlights significant correlations at p_spin_<0.05. Abbreviations: Attention Deficit Hyperactivity Disorder (ADHD), Autism Spectrum Disorder (ASD), Anorexia Nervosa (AN), Bipolar Disorder (BD), Major Depressive Disorder (MDD), Obsessive Compulsive Disorder (OCD), Schizophrenia (SCZ).

### Correlations Between GEDAR maps and Brain Structural Changes

Regional distributions of the GEDAR scores (Z-scores) are presented in Figures 3, 4 and 5 as calculated from both up-and down-regulated genes, only up-regulated and only down-regulated genes respectively. We display alongside maps of the regional distribution of ENIGMA effect sizes quantifying structural changes in patients compared to healthy controls for visual comparison.

**Figure 4.**
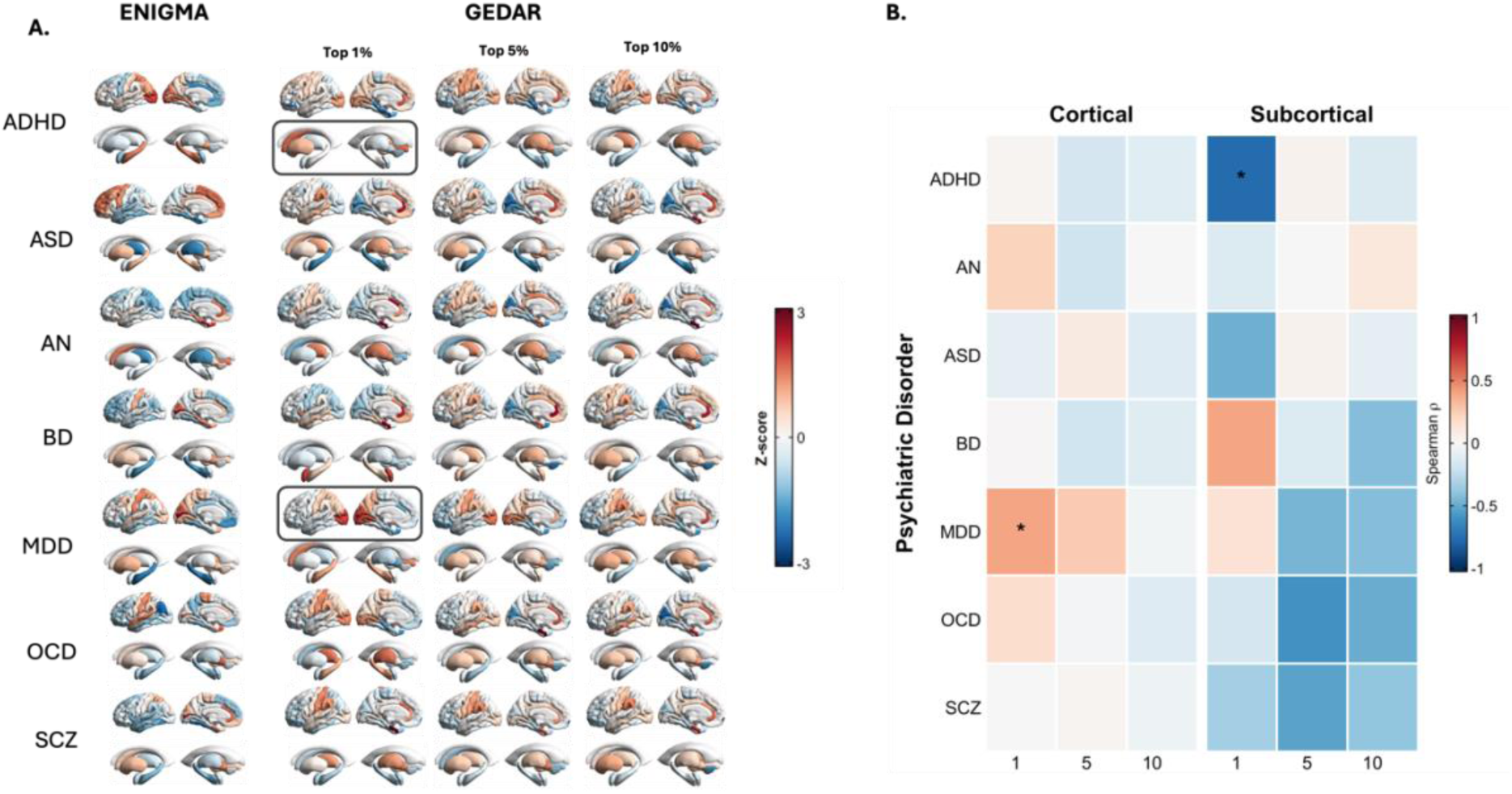
Correlation between ENIGMA and GEDAR brain maps (calculated from up-regulated TWAS genes). Panel A shows regional distributions of Z-scores of Cohen’s d effect sizes capturing changes in cortical thickness and subcortical volumes between patients and healthy-controls as provided by the respective ENIGMA meta-analysis (left panel); we also display our estimated GEDAR maps estimated from up-regulated TWAS genes by applying three different thresholds (top 10, 5 and 1%) to select the most differentially expressed genes. Panel B depicts spearman correlations between ENIGMA maps of structural differences and GEDAR maps calculated at the three different thresholds separately for cortical and subcortical regions. Significance was calculated by generating 1000 random spin permutations to account for spatial autocorrelation. The * highlights significant correlations at p_spin_<0.05. Abbreviations: Attention Deficit Hyperactivity Disorder (ADHD), Autism Spectrum Disorder (ASD), Anorexia Nervosa (AN), Bipolar Disorder (BD), Major Depressive Disorder (MDD), Obsessive Compulsive Disorder (OCD), Schizophrenia (SCZ).

**Figure 5.**
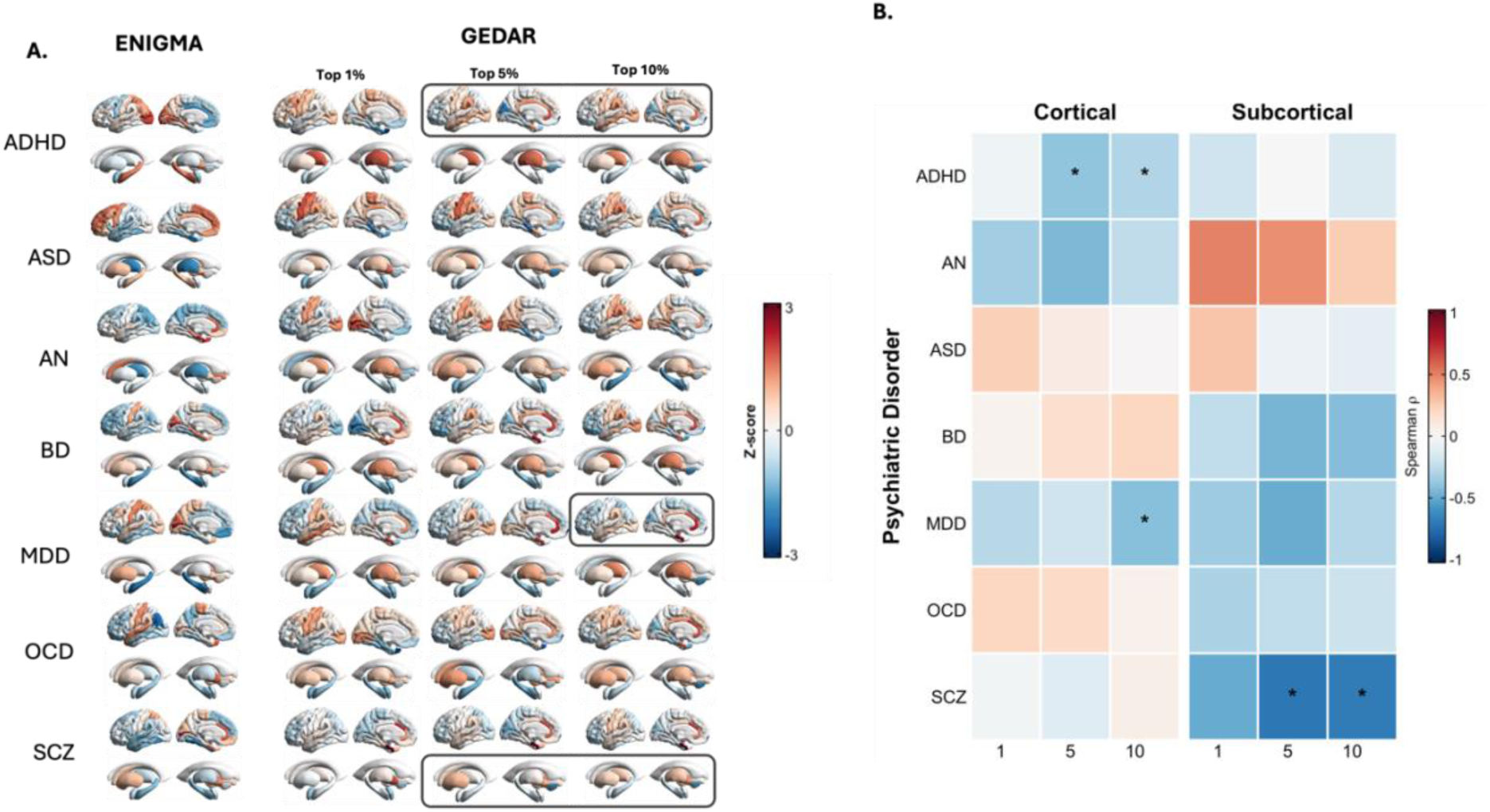
Correlation between ENIGMA and GEDAR brain maps (calculated from down-regulated TWAS genes). Panel A shows regional distributions of Z-scores of Cohen’s d effect sizes capturing changes in cortical thickness and subcortical volumes between patients and healthy-controls as provided by the respective ENIGMA meta-analysis (left panel); we also display our estimated GEDAR maps estimated from down-regulated TWAS genes by applying three different thresholds (top 10, 5 and 1%) to select the most differentially expressed genes. Panel B depicts spearman correlations between ENIGMA maps of structural differences and GEDAR maps calculated at the three different thresholds separately for cortical and subcortical regions. Significance was calculated by generating 1000 random spin permutations to account for spatial autocorrelation. The * highlights significant correlations at p_spin_<0.05. Abbreviations: Attention Deficit Hyperactivity Disorder (ADHD), Autism Spectrum Disorder (ASD), Anorexia Nervosa (AN), Bipolar Disorder (BD), Major Depressive Disorder (MDD), Obsessive Compulsive Disorder (OCD), Schizophrenia (SCZ).

#### Combined Up- and Down-Regulated Genes

In cortical regions, significant correlations were found for BD at the top 5% threshold (ρ = 0.456, *p_spin_* = 0.01), and for MDD at all thresholds (top 10%: ρ = 0.529, *p_spin_* = 0.005; top 5%: ρ = 0.546, *p_spin_* = 0.003; top 1%: ρ = 0.531, *p_spin_* = 0.003). In subcortical regions, significant correlations emerged for ADHD at the top 10% threshold (ρ = 0.919, *p_spin_* = 0.002), and for SCZ at both the top 10% (ρ = 0.703, *p_spin_* = 0.04) and 5% thresholds (ρ = –0.667, *p_spin_* = 0.04). None of the remaining correlations reached significance (Figure 2, Supplementary Table S1).

#### Up-Regulated Genes Only

In cortical regions, we found significant correlations for MDD at the top 1% threshold (ρ = 0.396, *p_spin_* = 0.031), while in subcortical regions only correlations for ADHD at the top 1% threshold reached significance (ρ = –0.775, *p_spin_* = 0.029). None of the remaining correlations reached significance (Figure 4, Supplementary Table S2).

#### Down-Regulated Genes Only

In cortical regions, we found significant correlations for ADHD at the top 10% (ρ = –0.300, *p_spin_* = 0.049) and 5% (ρ = –0.401, *p_spin_* = 0.009) thresholds, and for MDD at the top 10% threshold (ρ = –0.424, *p_spin_* = 0.017). In subcortical regions, SCZ showed significant negative correlations at both the top 10% (ρ = –0.703, *p_spin_* = 0.039) and 5% thresholds (ρ = –0.721, *p_spin_* = 0.04). None of the remaining correlations reached significance (Figure 5, Supplementary Table S3).

### GEDAR maps based on empirical differentially expressed genes in post-mortem brain samples of patients with psychiatric disorders

Correlation analyses using gene sets derived from meta-analytical investigations of differentially expressed genes determined in post-mortem brain samples yielded significant associations only for MDD. Significant correlations were found across all thresholds in both cortical (10%: ρ = –0.550, *p_spin_* = 0.002; 5%: ρ = –0.546, *p_spin_* = 0.002; 1%: ρ = –0.579, *p_spin_* < 0.001) and subcortical regions (10%, 5%, and 1% thresholds all ρ = –0.750, *p_spin_* ≈ 0.026– 0.029) suggesting that spatial patterns of gene expression alterations mirror regional brain structural changes in the disorder. No significant associations were found for ASD, BD, or SCZ (Figure 6).

**Figure 6.**
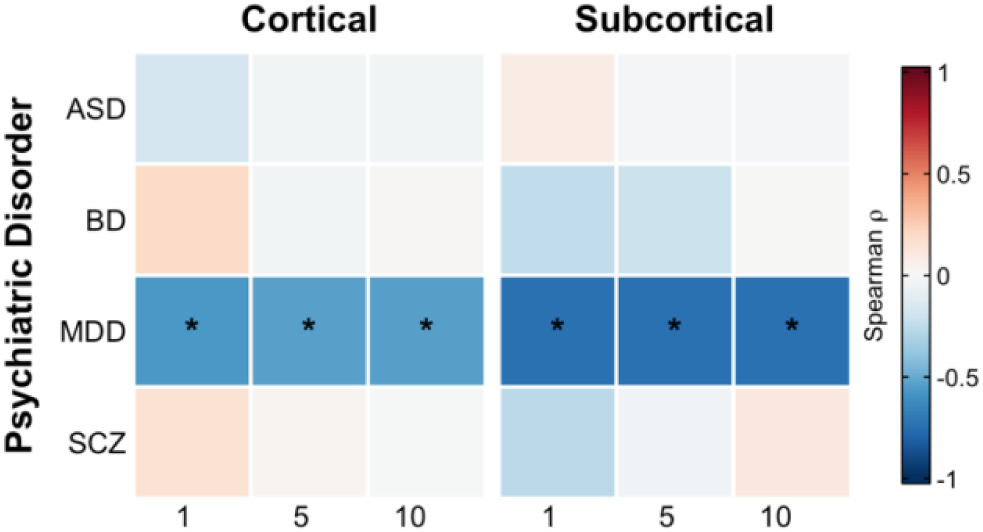
Correlation between ENIGMA and GEDAR brain maps (calculated from empirically determined differentially expressed genes). Spearman correlations between ENIGMA maps of structural changes and GEDAR maps calculated from differentially expressed genes in post-mortem brain samples at the three different thresholds separately for cortical and subcortical regions. Significance was calculated by generating 1000 random spin permutations to account for spatial autocorrelation. The * highlights significant correlations at p_spin_<0.05. Abbreviations: Attention Deficit Hyperactivity Disorder (ADHD), Autism Spectrum Disorder (ASD), Anorexia Nervosa (AN), Bipolar Disorder (BD), Major Depressive Disorder (MDD), Obsessive Compulsive Disorder (OCD), Schizophrenia (SCZ).

### Association between between-disorders heritability and GEDAR–ENIGMA spatial correlation

We computed Spearman correlations between heritability and GEDAR–ENIGMA spatial alignment across disorders for each combination of scheme (upregulated, downregulated, both), threshold (top 1, 5 and 10%), and brain compartment (cortical and subcortical). Correlation coefficients ranged from –0.929 to 0.607. Significant negative correlations (*p* < 0.05) were observed only for GEDARs calculated from up-regulated gene sets in cortical regions at the top 1% threshold, and in the combined gene sets in in the subcortical compartment at the top 1% threshold (Figure 7, Table 4).

**Figure 7.**
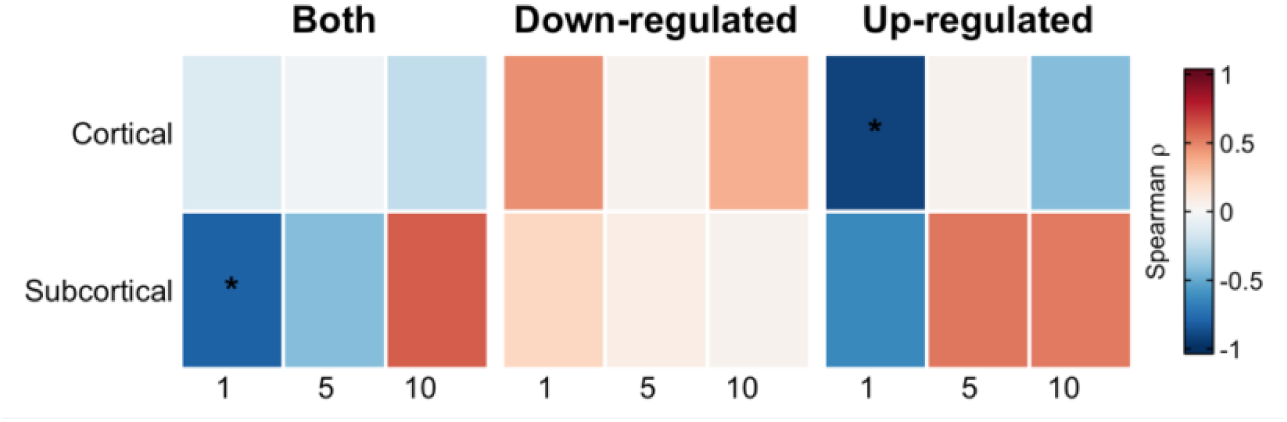
Correlations between heritability and GEDAR–ENIGMA correlation across disorders. The heatmaps depict Spearman correlations coefficients calculated between the middle point of heritability intervals and the estimated GEDAR-ENIGMA correlation. Both refers to GEDARs calculated from up- and down-regulated genes combined, while down- and up-regulated subplots refer to GEDARs calculate with either up- or down-regulated genes only. The * highlights significant correlations at a p<0.05.

**Table 4.**
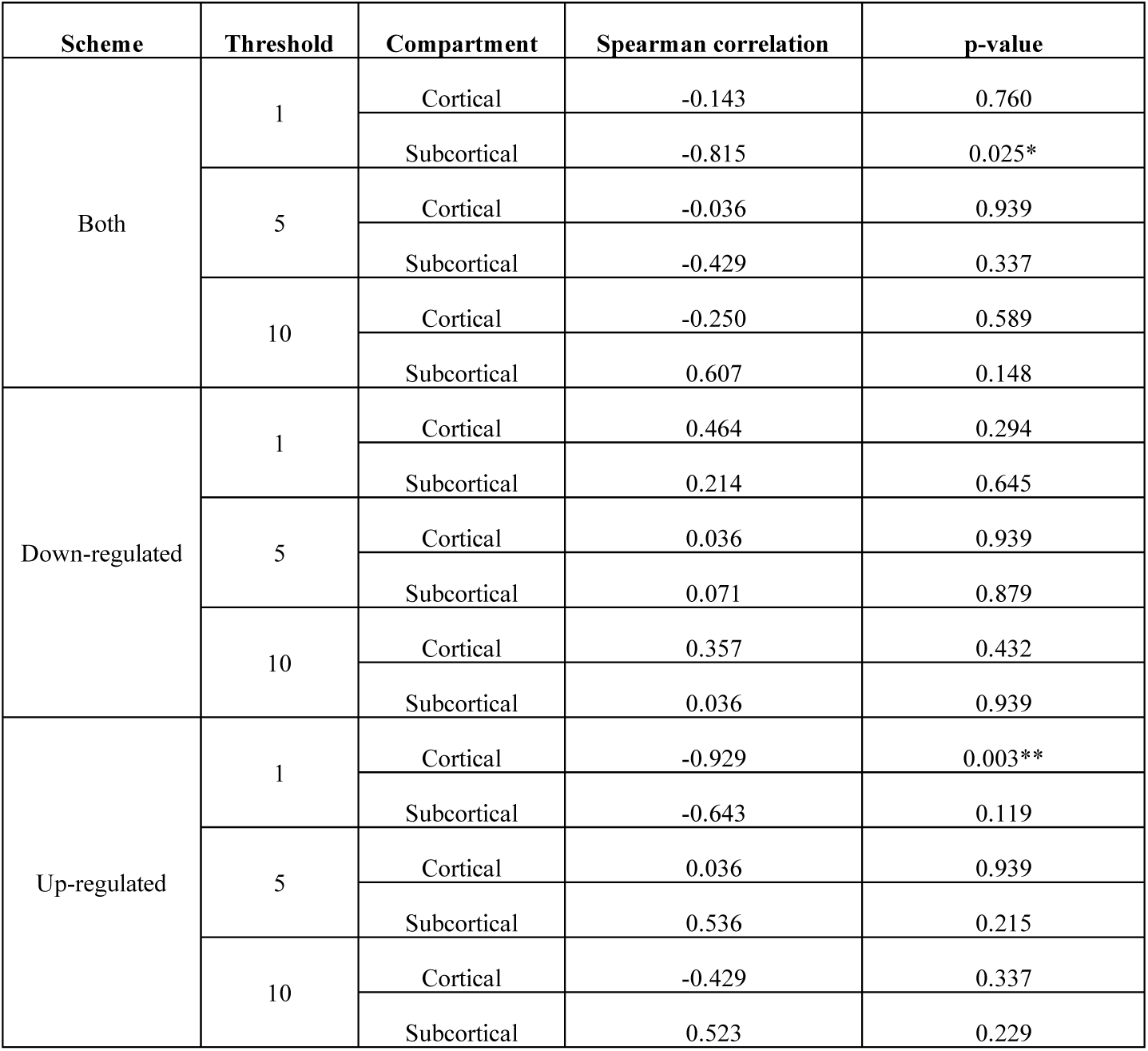
Correlations between heritability and GEDAR–ENIGMA correlation across disorders. Spearman correlations coefficients were calculated between the middle point of heritability intervals and the estimated GEDAR-ENIGMA correlation. Both refers to GEDARs calculated from up- and down-regulated genes combined, while down- and up-regulated subplots refer to GEDARs calculate with either up- or down-regulated genes only. The * highlights significant correlations at a p<0.05.

| Scheme | Threshold | Compartment | Spearman correlation | p-value |
| --- | --- | --- | --- | --- |
| Both | 1 | Cortical | -0.143 | 0.760 |
|  |  | Subcortical | -0.815 | 0.025* |
|  | 5 | Cortical | -0.036 | 0.939 |
|  |  | Subcortical | -0.429 | 0.337 |
|  | 10 | Cortical | -0.250 | 0.589 |
|  |  | Subcortical | 0.607 | 0.148 |
| Down-regulated | 1 | Cortical | 0.464 | 0.294 |
|  |  | Subcortical | 0.214 | 0.645 |
|  | 5 | Cortical | 0.036 | 0.939 |
|  |  | Subcortical | 0.071 | 0.879 |
|  | 10 | Cortical | 0.357 | 0.432 |
|  |  | Subcortical | 0.036 | 0.939 |
| Up-regulated | 1 | Cortical | -0.929 | 0.003** |
|  |  | Subcortical | -0.643 | 0.119 |
|  | 5 | Cortical | 0.036 | 0.939 |
|  |  | Subcortical | 0.536 | 0.215 |
|  | 10 | Cortical | -0.429 | 0.337 |
|  |  | Subcortical | 0.523 | 0.229 |

### Pathway Enrichment Analysis

#### Up-Regulated Pathways

For MDD, at the top 10% threshold, we found significant enrichment for two pathways involving **protein sequestering activity** (p_adj_ **=** 2.975 x 10^−2^) and the **MHC protein complex** (p_adj_ = 9.378 x 10^−3^) **(Supplementary Table S4)**.

#### Down-Regulated Pathways

In ADHD, at the top 10% threshold, we found significant enrichment for three pathways: **regulation of nervous system development** (p_adj_ **=** 1.585 x 10^−2^), **regulation of neurogenesis** (p_adj_ **=** 1.651 x 10^−2^), and **regulation of multicellular organismal development** (p_adj_ **=** 4.499 x 10^−2^). At the top 5% threshold, only enrichment for the **catenin complex** emerged as significant (p_adj_ **=** 2.011 x 10^−2^)—highlighting β-catenin’s role in cell adhesion (via cadherin– actin linkage) and Wnt signaling-mediated transcriptional regulation. For MDD, at the top 10% threshold, we found significant enrichment for 35 pathways. Several of these pathways were related to the **MHC complex** (e.g., protein complex binding p_adj_ **=** 8.406 x 10^−5^, class II components p_adj_ **=** 4.166 x 10^−2^), **immune response (e.g.,** p_adj_ **=** 6.290 x 10^−3^), **antigen processing (e.g.,** p_adj_ **=** 6.025 x 10^−4^), and broader **cellular components** (e.g., cell surface p_adj_ **=** 1.192 x 10^−4^, membrane side p_adj_ **=** 1.006 x 10^−3^, extracellular matrix p_adj_ **=** 3.754 x 10^−2^). For SCZ, at both the top 5 and 10% thresholds, we found significant enrichment for 53 pathways, including those involved in **immune activity** (MHC complex p_adj_ **=** 7.425 x 10^−3^, antigen binding p_adj_ **=** 1.029 x 10^−2^, adaptive immune response p_adj_ **=** 2.938 x 10^−3^), **cognition** (via MAPK/ERK-mediated synaptic plasticity p_adj_ **=** 3.070 x 10^−2^), and **leukocyte regulation** (involving JAK-STAT and NF-κB pathways p_adj_ **=** 1.799 x 10^−2^) **(Supplementary Table S1).**

## Discussion

This study introduces a novel and generalizable framework for mapping polygenic transcriptional risk onto macroscale brain phenotypes in psychiatric disorders. By integrating genetically predicted gene expression profiles from transcriptome-wide association studies (TWAS) with spatial transcriptomic data from the Allen Human Brain Atlas and structural neuroimaging meta-analyses from the ENIGMA consortium, we sought to clarify how polygenic risk translates into regional brain structural alterations. Our findings demonstrate that neuroimaging phenotypes in certain disorders reflect underlying, spatially patterned transcriptional dysregulation linked to genetic liability. This integrative approach not only advances mechanistic understanding of psychiatric disease but also lays the groundwork for translational strategies that connect genome, transcriptome, and brain structure.

We found that predicted transcriptomic dysregulation captured by GEDAR scores significantly correlated with regional brain structural differences with HCs in a subset of disorders. These effects were strongest in MDD, where robust associations were observed across cortical and subcortical regions and across multiple thresholds of gene inclusion. Notably, both TWAS-predicted and empirically-derived differentially expressed genes from post-mortem brain samples yielded significant correlations in MDD, reinforcing the idea that transcriptomic signatures—whether genetically inferred or empirically derived—are spatially aligned with the disorder’s neuroanatomical profile. These findings suggest that MDD-related structural changes, which often involve limbic and prefrontal cortices, may partly reflect regionally patterned gene expression dysregulation. Pathway analyses revealed that many of the enriched biological processes in MDD were immune-related, including components of the major histocompatibility complex (MHC). These findings are consistent with prior evidence linking immune dysregulation, cortical thinning, and associated brain structural changes in MDD^51–54^. The enrichment of these pathways highlights not only the regional impacts of transcriptomic dysregulation on brain anatomy, but also the critical role of immune and inflammatory signalling in the pathophysiology of MDD^52,55,56^. Such insights may help inform therapeutic strategies, including current efforts to target immune-related pathways^57^ to mitigate the structural and functional consequences of the disorder. More broadly, these findings underscore the importance of integrating genetic and transcriptomic data with brain imaging to uncover biologically grounded mechanisms driving psychiatric disorders.

In contrast, other disorders such as schizophrenia (SCZ) and attention-deficit/hyperactivity disorder (ADHD) exhibited significant associations primarily in subcortical regions. For SCZ, these were predominantly driven by down-regulated genes and involved key immune-related transcripts, including several MHC-related genes, highlighting the role of neuroimmune interactions in subcortical vulnerability^58–60^. In ADHD, gene expression changes aligned with known striatal alterations and were enriched for pathways involved in neurodevelopment and cell adhesion, such as the catenin complex and Wnt signalling^36,61–63^. The catenin complex, in particular, is crucial for establishing and maintaining neuronal connectivity during brain development, as it mediates cell–cell adhesion through interactions with cadherins at synaptic junctions. β-catenin, a key component of this complex, also acts as an intracellular signalling molecule downstream of Wnt, influencing gene expression patterns important for neuronal proliferation, differentiation, and synaptic plasticity ^64,65^. These associations highlight how transcriptomic perturbations may contribute to regional specificity in structural brain differences observed in ADHD, reinforcing the idea that subcortical circuits, especially within the striatum, may be uniquely sensitive to genetically regulated developmental processes in early-onset psychiatric disorders^66,67^. These insights further support existing neurodevelopmental frameworks and pinpoint the catenin complex as a biologically plausible target for future mechanistic studies and therapeutic interventions in ADHD and related neurodevelopmental conditions.

We note, however, that methodological choices regarding thresholds for gene inclusion in GEDAR score calculations appeared influential in specific cases. For example, in schizophrenia, GEDAR scores derived from combined sets of up- and down-regulated genes at a less stringent threshold (top 5%) resulted in a shift in the correlation direction from positive to negative compared to the stricter threshold (top 1%). When analyses were conducted separately for up- and down-regulated genes, negative associations generally emerged, suggesting potential opposing transcriptional effects. While we included combined up- and down-regulated gene sets to maximize sensitivity to spatial associations, these observations indicate that interpreting combined transcriptional signatures warrants caution. Specifically, Transcriptome-Wide Association Study (TWAS) Z-scores represent associations with a given trait rather than direct evidence of gene regulation, and combining genes with opposing expression directions inherently assumes shared directionality of biological effects, potentially obscuring true regional expression patterns^25^.

Interestingly, bipolar disorder (BD) showed more modest correspondence between GEDAR and neuroimaging phenotypes, with significant effects emerging only at specific thresholds. For other disorders, including autism spectrum disorder (ASD), anorexia nervosa (AN), and obsessive-compulsive disorder (OCD), associations were limited or absent. This variability most likely reflect the extent to which observed brain changes are mediated by factors not captured by genetically predicted expression—such as post-transcriptional regulation^17^, alternative splicing^68–70^, epigenetic modifications^71^, environmental exposures^72^ or developmental timing^73^. To clarify whether the limited predictive power in some disorders was due to the use of TWAS-predicted genes rather than actual transcriptional abnormalities, we also tested GEDAR maps constructed from empirically identified differentially expressed genes. These analyses again yielded significant associations only for MDD, suggesting that in other disorders, the spatial distribution of gene expression—regardless of whether inferred or observed—may not align with the anatomical profile of structural alterations. This points to a fundamental challenge: while gene expression is undoubtedly relevant to brain structure and function, its contribution may be more salient in some disorders than others or may operate through intermediate biological mechanisms not readily captured by bulk transcriptomic data.

The differential strength of transcriptomic–anatomical associations across psychiatric disorders is intriguing in itself and might provide important biological insights. While disorders like MDD demonstrated robust associations, the limited or absent correlations observed in ASD, AN, and OCD, which predominantly fall into neurodevelopmental categories^74^, may reflect critical differences in developmental timing rather than methodological shortcomings. Specifically, the transcriptomic signatures most relevant to neurodevelopmental disorders likely occur during prenatal or early-life periods^75,76^, long before structural alterations become evident on adult MRI scans. Because TWAS predominantly capture genetically regulated, relatively stable patterns of gene expression based primarily on adult reference data, these approaches align closely with conditions like MDD, characterized by ongoing pathological processes in adulthood. Conversely, for disorders such as ASD or AN, early developmental mechanisms may be paramount and thus not adequately captured by adult-derived transcriptomic maps. Framing our results in this context provides a meaningful interpretation and underscores the need for future research incorporating developmentally informed gene expression datasets and longitudinal imaging–transcriptomic designs to better capture the critical windows during which gene expression influences neuroanatomical development. However, another relevant consideration is the potential confounding effect of psychotropic medication use in ENIGMA cohorts, particularly in disorders such as schizophrenia, bipolar disorder, and major depression. While our transcriptomic predictions are genetically driven, it remains possible that medication exposure could shape regional structural phenotypes in ways that amplify—or obscure—GEDAR alignment. This may partly explain the absence of significant correlations in less medicated cohorts such as ASD or AN. Future analyses stratifying patients by medication status or illness duration (e.g., first-episode vs chronic) could help clarify this issue.

Although we observed some tendency for disorders with higher heritability, such as SCZ and ADHD, to show stronger GEDAR–neuroimaging correspondence, this relationship was not uniform. MDD, which has moderate heritability, yielded some of the most consistent associations, while BD and ASD—with higher heritability—showed weaker to no effects. Further reinforcing this interpretation, we could not find a possible positive association between heritability and the strength of spatial association between GEDAR and ENIGMA maps of structural changes. Where this reached significance, the association was in fact negative, which contradicts our initial predictions. These findings suggest that heritability alone does not determine how well gene expression predicts brain changes; rather, the molecular architecture, developmental profile, and pathophysiological mechanisms specific to each disorder likely modulate this relationship^72,77^, highlighting that further research is needed to clarify the precise neurobiological mechanisms underlying changes in brain structure documented in patients with psychiatric disorders through the use of neuroimaging techniques.

Several limitations should be acknowledged. First, GEDAR maps are based on predicted transcriptomic alterations and may not fully capture the complexity of gene regulation in the human brain^78^. Second, while the AHBA provides valuable spatial resolution, its donor sample is small and may not generalize across populations^2,14^. Third, the reliance on summary-level imaging and genetic data limits our ability to assess individual-level variation when psychiatric disorders and associated structural brain changes are now understood to be highly heterogenous at a regional level^79–81^. Fourth, it is also important to note that our TWAS approach involved averaging across all S-PrediXcan tissue panels, rather than tailoring gene expression prediction to specific brain regions. While using distinct tissue panels for each brain region could potentially offer more anatomical specificity, this strategy is currently constrained by the lack of direct correspondence between the GTEx panels and the Desikan–Killiany atlas. Some GTEx tissues map cleanly to multiple brain regions, while others map ambiguously or not at all. To address this, we prioritized anatomical coverage over regional specificity, trading a degree of transcriptomic granularity for the ability to analyse all available ENIGMA regions uniformly. Future work could explore hybrid approaches that combine region-specific predictions where feasible or integrate emerging resources that offer finer-grained transcriptomic spatial mapping. Fifth, our analyses used adult-derived transcriptomic reference models (GTEx v8 and AHBA), which may not capture developmentally critical transcriptional patterns relevant to early-onset disorders. The release of neurodevelopmentally informed eQTL resources (e.g., the human and non-human developmental GTex^82^) will allow future studies to more accurately model the spatiotemporal dynamics of gene expression during prenatal and early postnatal development, a critical period for disorders such as ASD, ADHD, and AN.

Overall, this study provides preliminary evidence supporting the notion that genetically driven transcriptomic dysregulation is spatially aligned, to varying degrees, with regional brain structural changes in some psychiatric disorders, particularly MDD. Our findings highlight the potential of this integrative framework to advance our understanding of the molecular basis of disease-related brain changes while underscoring the need for refined, multi-omic, and cell-type-specific methods to capture the full spectrum of gene expression dynamics^83,84^. As our understanding of the molecular and cellular complexity of the human brain expands, future research should aim to incorporate additional biological layers and multi-omics data, such as single-cell transcriptomics, alternative splicing and epigenetics to build a more comprehensive understanding of how genetically encoded molecular changes might map on the full cascade from genome to brain to behaviour and shape the neuroanatomy of psychiatric disorders.

## Acknowledgments

This study is funded by the National Institute for Health and Care Research (NIHR) Maudsley Biomedical Research Centre, South London and Maudsley NHS Trust. The views expressed are those of the author(s) and not necessarily those of the NIHR or the Department of Health and Social Care. DM is supported by NIHR Maudsley Biomedical Research Centre, South London and Maudsley NHS Trust. DD and FT are partially supported by NIHR Maudsley Biomedical Research Centre, South London and Maudsley NHS Trust. T.R.P. is supported by an MRC (UKRI) New Investigator Research Grant (MR/W028018/1). For the purpose of open access, the author has applied a CC BY public copyright license to any Author Accepted Manuscript version arising from this submission. This research is also part-funded by the National Institute for Health and Care Research (NIHR) Maudsley Biomedical Research Centre. The views expressed are those of the authors and not necessarily those of the NHS, the NIHR or the Department of Health and Social Care. R.R.R.D. and T.R.P. are supported by a Psychiatry Research Trust Grant.

## CreDit statement

AG - data curation, formal analysis, writing (original draft), writing (review and editing).

TP – conceptualisation, writing (review and editing)

RD – conceptualisation, writing (review and editing).

GN - data curation, formal analysis, writing (review and editing).

FT - writing (review and editing), funding acquisition.

SCRW - writing (review and editing), funding acquisition.

MV - writing (review and editing)

DM – conceptualisation, writing (original draft), writing (review and editing), supervision (shared).

DD – conceptualisation, writing (original draft), writing (review and editing), supervision (shared).

