## Supplementary Materials for "Transcriptome-informed brain cartography of polygenic risk and association with brain structure in major psychiatric disorders"

[1] *Department of Neuroimaging, Institute of Psychiatry, Psychology and Neuroscience, King's College London, London, UK*
[2] *Department of Child and Adolescent Psychiatry, University Hospital, Goethe University, Frankfurt am Main, Germany*
[3] *Cooperative Brain Imaging Centre (COBIC), Goethe University, Frankfurt am Main, Germany*
[4] *Social, Genetic & Developmental Psychiatry Centre, Institute of Psychiatry, Psychology & Neuroscience, King's College London, London, UK*
[5] *The Institute for Human and Synthetic Minds, King's College London, London, UK*
[6] *Department of Information Engineering, University of Padova, Padova, Italy*
[7] *Department of Psychology, City St George's, University of London, London, UK*
[*] *Equal contribution*

### Preliminary analyses with H-MAGMA

To develop a spatial model of regional variation of structural brain differences from transcriptomic architecture the first step is to convert polygenic risk from the genetic variant space to the gene level space. To achieve this, we used an algorithm of risk calculation at the level of gene such as Multi-marker Analysis of GenoMic Association (MAGMA) (de Leeuw et al., 2015).

MAGMA is a widely used tool for gene-based and gene-set association analyses in genome-wide association studies (GWAS). It aggregates SNP-level association statistics into gene-level p-values by applying a multiple linear regression framework that accounts for linkage disequilibrium (LD) and gene size. In our case we performed the analysis using a refined version of MAGMA that is well suited for brain tissues, namely H-MAGMA (Sey et al., 2020). Unlike traditional MAGMA, which assigns SNPs to genes based on linear proximity, H-MAGMA uses Hi-C chromatin interaction maps to map non-coding variants to their regulatory target genes. This resulted in a list of prioritised genes based on predicted cumulative risk of dysregulation.

We then developed a dysregulation score for each region of the Desikan-Killiany (DK) atlas by combining the regional gene expression pattern, as expressed by the Allen Human Brain

Atlas (AHBA), with the weight indicated by H-MAGMA in those genes that showed p-value<0.001. Finally, we correlated these scores with meta-analytical maps of structural brain abnormalities measured by cortical thickness of six psychiatric disorders (ADHD, AN, ASD, BD, MDD, and SCZ) from the Enhancing Imaging Genetics through Meta Analysis (ENIGMA) initiative.

With this approach only one MDD showed significant correlation between H-MAGMA identified genes in the brain and ENIGMA measured regional abnormalities ($\rho=0.346$, $p_{spin}=0.042$). This approach nonetheless lacked the information about the directionality of the changes. Directionality might play a significant role in psychiatric disorders as some changes might be a protective factor while other might be pathological.

**Supplementary Figure 1. Sampling sites for GTEx prediction models**. The Genotype-Tissue Expression (GTEx) project is a large-scale initiative that characterizes the relationship between genetic variation and gene expression across multiple human tissues. By integrating genomic and transcriptomic data from post-mortem donors, GTEx provides a comprehensive resource for studying tissue-specific gene regulation and identifying expression quantitative trait loci (eQTLs). For brain-related research, GTEx includes RNA-sequencing data from multiple brain regions, such as the cortex, cerebellum, hippocampus, basal ganglia (including the caudate, putamen, and nucleus Accumbens), hypothalamus, amygdala, and substantia nigra. These regions are critical for understanding neurodevelopment, cognition, and neuropsychiatric disorders. GTEx-based elastic net models use cis-genetic variation (within 1 Mb of a gene) to predict gene expression in specific tissues. Elastic net regression combines lasso (L1) and ridge (L2) penalties to select relevant genetic variants while maintaining model stability. In the present study we used the models here described to predict the transcriptomic profile from GWAS data in each tissue and then combine them for a whole brain approach. More information about the GTEx, GTEx panels and methods are available at https://www.gtexportal.org/home/.

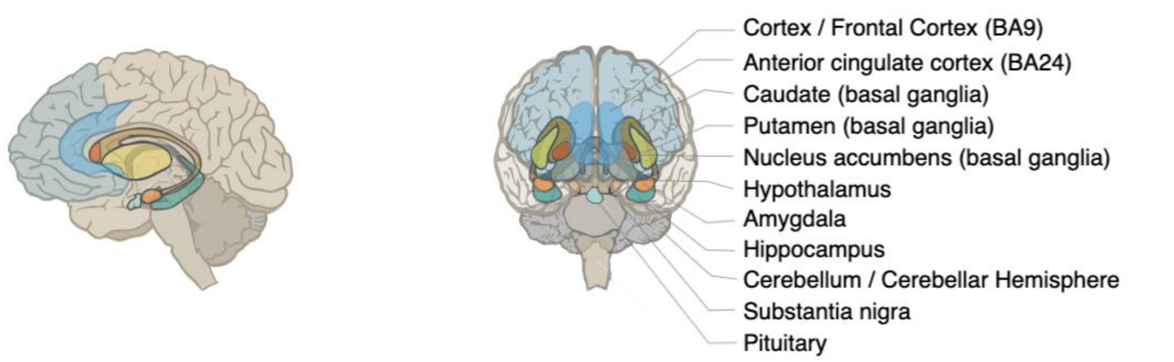


**Supplementary Table S1.** The table shows results for correlation analysis between GEDAR brain maps and brain structural changes. Table shows GEDAR maps for combined up- and down-regulated genes. The * indicates significant p<0.05.

| Disorder | Threshold | Compartment | Spearman correlation | p-value |
|---|---|---|---|---|
| ADHD | 1 | Cortical | 0.130 | 0.274 |
| | | Subcortical | -0.396 | 0.181 |
| | 5 | Cortical | 0.151 | 0.273 |
| | | Subcortical | 0.054 | 0.470 |
| | 10 | Cortical | 0.252 | 0.131 |
| | | Subcortical | 0.920 | 0.002* |
| AN | 1 | Cortical | -0.517 | 0.056 |
| | | Subcortical | 0.214 | 0.349 |
| | 5 | Cortical | -0.437 | 0.098 |
| | | Subcortical | 0.321 | 0.240 |
| | 10 | Cortical | -0.408 | 0.122 |
| | | Subcortical | 0.071 | 0.418 |
| ASD | 1 | Cortical | -0.150 | 0.252 |
| | | Subcortical | -0.262 | 0.289 |
| | 5 | Cortical | 0.413 | 0.073 |
| | | Subcortical | 0.206 | 0.316 |
| | 10 | Cortical | -0.053 | 0.423 |
| | | Subcortical | 0.449 | 0.154 |
| BD | 1 | Cortical | 0.175 | 0.153 |
| | | Subcortical | 0.214 | 0.303 |
| | 5 | Cortical | 0.456 | 0.01** |
| | | Subcortical | -0.321 | 0.229 |
| | 10 | Cortical | 0.314 | 0.09 |
| | | Subcortical | -0.571 | 0.089 |
| MDD | 1 | Cortical | 0.531 | 0.003** |
| | | Subcortical | 0.571 | 0.083 |
| | 5 | Cortical | 0.547 | 0.003** |
| | | Subcortical | 0.571 | 0.091 |
| | 10 | Cortical | 0.529 | 0.005** |
| | | Subcortical | -0.143 | 0.392 |
| OCD | 1 | Cortical | -0.191 | 0.111 |
| | | Subcortical | 0.214 | 0.311 |
| | 5 | Cortical | -0.097 | 0.266 |
| | | Subcortical | -0.250 | 0.283 |
| | 10 | Cortical | -0.211 | 0.086 |
| | | Subcortical | -0.429 | 0.161 |
| SCZ | 1 | Cortical | -0.101 | 0.331 |
| | | Subcortical | -0.288 | 0.242 |
| | 5 | Cortical | -0.002 | 0.486 |

| | | | | |
|---|---|---|---|---|
| | | Subcortical | -0.667 | 0.047* |
| | 10 | Cortical | -0.333 | 0.057 |
| | | Subcortical | 0.702 | 0.041* |

**Supplementary Table S2.** The table shows results for correlation analysis between GEDAR brain maps and brain structural changes. Table shows GEDAR maps for up-regulated genes. The * indicates significant p<0.05.

| Disorder | Threshold | Compartment | Spearman correlation | p-value |
|---|---|---|---|---|
| ADHD | 1 | Cortical | 0.024 | 0.458 |
| | | Subcortical | -0.775 | 0.029* |
| | 5 | Cortical | -0.178 | 0.235 |
| | | Subcortical | 0.036 | 0.484 |
| | 10 | Cortical | -0.116 | 0.273 |
| | | Subcortical | -0.144 | 0.347 |
| AN | 1 | Cortical | 0.223 | 0.125 |
| | | Subcortical | -0.143 | 0.375 |
| | 5 | Cortical | -0.202 | 0.166 |
| | | Subcortical | 0.000 | 0.468 |
| | 10 | Cortical | -0.012 | 0.512 |
| | | Subcortical | 0.107 | 0.422 |
| ASD | 1 | Cortical | -0.095 | 0.3-5 |
| | | Subcortical | -0.486 | 0.131 |
| | 5 | Cortical | 0.092 | 0.340 |
| | | Subcortical | 0.037 | 0.476 |
| | 10 | Cortical | -0.093 | 0.416 |
| | | Subcortical | 0.449 | 0.154 |
| BD | 1 | Cortical | 0.001 | 0.524 |
| | | Subcortical | 0.393 | 0.184 |
| | 5 | Cortical | -0.195 | 0.142 |
| | | Subcortical | -0.143 | 0.376 |
| | 10 | Cortical | -0.126 | 0.247 |
| | | Subcortical | -0.428 | 0.163 |
| MDD | 1 | Cortical | 0.396 | 0.031* |
| | | Subcortical | 0.143 | 0.376 |
| | 5 | Cortical | 0.257 | 0.131 |
| | | Subcortical | -0.464 | 0.135 |
| | 10 | Cortical | -0.028 | 0.404 |
| | | Subcortical | -0.428 | 0.181 |
| OCD | 1 | Cortical | 0.168 | 0.125 |

| | | | | |
|---|---|---|---|---|
| | | Subcortical | -0.178 | 0.321 |
| | 5 | Cortical | -0.025 | 0.440 |
| | | Subcortical | -0.607 | 0.060 |
| | 10 | Cortical | -0.128 | 0.233 |
| | | Subcortical | -0.500 | 0.124 |
| SCZ | 1 | Cortical | -0.009 | 0.490 |
| | | Subcortical | -0.342 | 0.212 |
| | 5 | Cortical | 0.025 | 0.438 |
| | | Subcortical | -0.541 | 0.101 |
| | 10 | Cortical | -0.05 | 0.360 |
| | | Subcortical | -0.396 | 0.194 |

**Supplementary Table S3.** The table shows results for correlation analysis between GEDAR brain maps and brain structural changes. Table shows GEDAR maps for down-regulated genes. The * indicates significant p<0.05.

| Disorder | Threshold | Compartment | Spearman correlation | p-value |
|---|---|---|---|---|
| ADHD | 1 | Cortical | -0.045 | 0.412 |
| | | Subcortical | -0.198 | 0.319 |
| | 5 | Cortical | -0.401 | 0.009** |
| | | Subcortical | 0.000 | 0.476 |
| | 10 | Cortical | -0.300 | 0.049* |
| | | Subcortical | -0.144 | 0.347 |
| AN | 1 | Cortical | -0.348 | 0.124 |
| | | Subcortical | 0.214 | 0.349 |
| | 5 | Cortical | -0.452 | 0.083 |
| | | Subcortical | 0.321 | 0.240 |
| | 10 | Cortical | -0.252 | 0.211 |
| | | Subcortical | 0.071 | 0.418 |
| ASD | 1 | Cortical | 0.238 | 0.316 |
| | | Subcortical | 0.280 | 0.271 |
| | 5 | Cortical | 0.082 | 0.411 |
| | | Subcortical | -0.056 | 0.442 |
| | 10 | Cortical | 0.001 | 0.498 |
| | | Subcortical | -0.093 | 0.425 |
| BD | 1 | Cortical | 0.033 | 0.439 |
| | | Subcortical | -0.250 | 0.287 |
| | 5 | Cortical | 0.168 | 0.185 |
| | | Subcortical | -0.464 | 0.133 |
| | 10 | Cortical | 0.209 | 0.132 |

| | | | | |
|---|---|---|---|---|
| | | Subcortical | -0.428 | 0.162 |
| MDD | 1 | Cortical | -0.278 | 0.066 |
| | | Subcortical | -0.357 | 0.214 |
| | 5 | Cortical | -0.201 | 0.144 |
| | | Subcortical | -0.500 | 0.131 |
| | 10 | Cortical | -0.424 | 0.017 |
| | | Subcortical | -0.286 | 0.249 |
| OCD | 1 | Cortical | 0.207 | 0.088 |
| | | Subcortical | -0.321 | 0.240 |
| | 5 | Cortical | 0.189 | 0.125 |
| | | Subcortical | -0.250 | 0.288 |
| | 10 | Cortical | 0.046 | 0.408 |
| | | Subcortical | -0.214 | 0.326 |
| SCZ | 1 | Cortical | -0.032 | 0.445 |
| | | Subcortical | -0.505 | 0.130 |
| | 5 | Cortical | -0.120 | 0.246 |
| | | Subcortical | -0.721 | 0.040* |
| | 10 | Cortical | 0.059 | 0.365 |
| | | Subcortical | -0.703 | 0.039* |

**Supplementary Table S4.** The table shows significantly enriched pathways emerging from the pathway enrichment analysis on the lists of genetically-predicted differentially regulated genes from TWAS for those disorders for which we found a significant association between GEDAR and ENIGMA maps.

| Disorder | Threshold (%) | Gene Regulation | Geneset Source | Term ID | Term Name | $p_{adj}$ |
|---|---|---|---|---|---|---|
| MDD | 1 | UP | GO:MF | GO:0140311 | Protein sequestering activity | $2.975 \times 10^{-2}$ |
| | | | GO:CC | GO:0042611 | MHC protein complex | $9.378 \times 10^{-3}$ |
| ADHD | 10 | DOWN | GO:BP | GO: 0051960 | Regulation of nervous system development | $1.585 \times 10^{-2}$ |
| | | | GO:BP | GO: 0050767 | Regulation of neurogenesis | $1.651 \times 10^{-2}$ |
| | | | GO:BP | GO: 0051960 | Regulation of multicellular organismal development | $4.499 \times 10^{-2}$ |
| | 5 | DOWN | GO:CC | GO: 0016342 | Catenin complex | $2.011 \times 10^{-2}$ |
| MDD | 10 | DOWN | GO:CC | GO:0042613 | MHC class II protein | $4.166 \times 10^{-2}$ |

| | | | | | | |
|---|---|---|---|---|---|---|
| | | | | | complex | |
| | | | GO:CC | GO:0042611 | MHC protein complex | $8.406 \times 10^{-5}$ |
| | | | GO:MF | GO:0023023 | MHC protein complex binding | $3.242 \times 10^{-2}$ |
| | | | GO:MF | GO:0003823 | Antigen binding | $1.053 \times 10^{-2}$ |
| | | | GO:BP | GO:0019882 | antigen processing and presentation | $6.025 \times 10^{-4}$ |
| | | | GO:BP | GO:0019883 | antigen processing and presentation of endogenous antigen | $4.860 \times 10^{-2}$ |
| | | | GO:BP | GO:0002483 | antigen processing and presentation of endogenous peptide antigen | $1.693 \times 10^{-2}$ |
| | | | GO:BP | GO:0019884 | antigen processing and presentation of exogenous antigen | $1.226 \times 10^{-2}$ |
| | | | GO:BP | GO:0019886 | antigen processing and presentation of exogenous peptide antigen via MHC class II | $3.205 \times 10^{-2}$ |
| | | | GO:BP | GO:0048002 | antigen processing and presentation of peptide antigen | $2.126 \times 10^{-4}$ |
| | | | GO:BP | GO:0007155 | cell adhesion | $2.523 \times 10^{-2}$ |
| | | | GO:CC | GO:0071944 | cell periphery | $1.216 \times 10^{-6}$ |
| | | | GO:CC | GO:0009986 | cell surface | $1.192 \times 10^{-4}$ |
| | | | GO:CC | GO:0062023 | collagen-containing extracellular matrix | $1.496 \times 10^{-2}$ |

| | | | | | | |
|---|---|---|---|---|---|---|
| | | | GO:CC | GO:0030312 | external encapsulating structure | $3.817 \times 10^{-2}$ |
| | | | GO:CC | GO:0009897 | external side of plasma membrane | $1.006 \times 10^{-3}$ |
| | | | GO:CC | GO:0031012 | extracellular matrix | $3.754 \times 10^{-2}$ |
| | | | GO:BP | GO:0002768 | immune response-regulating cell surface receptor signaling pathway | $3.982 \times 10^{-2}$ |
| | | | GO:CC | GO:0098553 | lumenal side of endoplasmic reticulum membrane | $4.329 \times 10^{-3}$ |
| | | | GO:CC | GO:0098576 | lumenal side of membrane | $1.917 \times 10^{-2}$ |
| | | | GO:CC | GO:0016020 | membrane | $3.789 \times 10^{-4}$ |
| | | | GO:MF | GO:0060089 | molecular transducer activity | $1.764 \times 10^{-2}$ |
| | | | GO:MF | GO:0042605 | peptide antigen binding | $1.583 \times 10^{-4}$ |
| | | | GO:MF | GO:0042277 | peptide binding | $2.378 \times 10^{-2}$ |
| | | | GO:CC | GO:0005886 | plasma membrane | $4.688 \times 10^{-5}$ |
| | | | GO:BP | GO:1902533 | positive regulation of intracellular signal transduction | $3.090 \times 10^{-2}$ |

| | | | | | | |
|---|---|---|---|---|---|---|
| | | | GO:BP | GO:0048584 | positive regulation of response to stimulus | $2.011 \times 10^{-2}$ |
| | | | GO:BP | GO:0097104 | postsynaptic membrane assembly | $3.615 \times 10^{-2}$ |
| | | | GO:BP | GO:0002822 | regulation of adaptive immune response based on somatic recombination of immune receptors built from immunoglobulin superfamily domains | $2.325 \times 10^{-2}$ |
| | | | GO:BP | GO:0050770 | regulation of axonogenesis | $6.242 \times 10^{-3}$ |
| | | | GO:BP | GO:0050776 | regulation of immune response | $6.290 \times 10^{-3}$ |
| | | | GO:BP | GO:0048583 | regulation of response to stimulus | $4.235 \times 10^{-2}$ |
| | | | GO:CC | GO:0098552 | side of membrane | $8.083 \times 10^{-3}$ |
| | | | GO:MF | GO:0038023 | signaling receptor activity | $1.764 \times 10^{-2}$ |
| SCZ | 10 | DOWN | GO:MF | GO:0042605 | peptide antigen binding | $3.65 \times 10^{-6}$ |
| | 5 | | | | | $2.60 \times 10^{-6}$ |
| | 10 | | GO:MF | GO:0046977 | TAP binding | $3.999 \times 10^{-3}$ |

| | | | | | | |
|---|---|---|---|---|---|---|
| | 10 | | GO:MF | GO:0042277 | peptide binding | $1.657 \times 10^{-2}$ |
| | 5 | | | | | $8.399 \times 10^{-3}$ |
| | 10 | | GO:MF | GO:0003823 | antigen binding | $5.957 \times 10^{-2}$ |
| | 5 | | | | | $1.029 \times 10^{-2}$ |
| | 5 | | GO:MF | GO:0023026 | MHC class II protein complex binding | $3.000 \times 10^{-2}$ |
| | 10 | | GO:BP | GO:0048002 | antigen processing and presentation of peptide antigen | $3.174 \times 10^{-7}$ |
| | 5 | | | | | $2.220 \times 10^{-6}$ |
| | 10 | | GO:BP | GO:0019882 | antigen processing and presentation | $3.898 \times 10^{-5}$ |
| | 5 | | | | | $9.565 \times 10^{-5}$ |
| | 10 | | GO:BP | GO:0002476 | antigen processing and presentation of endogenous peptide antigen via MHC class Ib | $2.264 \times 10^{-3}$ |
| | 5 | | | | | $2.085 \times 10^{-2}$ |
| | 10 | | GO:BP | GO:0002484 | antigen processing and presentation of endogenous peptide antigen via MHC class I via ER pathway | $2.264 \times 10^{-3}$ |
| | 5 | | | | | $2.085 \times 10^{-2}$ |
| | 10 | | GO:BP | GO:0019886 | antigen processing and presentation of exogenous peptide antigen via MHC class II | $3.205 \times 10^{-2}$ |
| | 5 | | | | | $2.563 \times 10^{-3}$ |

| | | | | | | |
|---|---|---|---|---|---|---|
| | 10 | | GO:BP | GO:0002822 | regulation of adaptive immune response based on somatic recombination of immune receptors built from immunoglobulin superfamily domains | $2.325 \times 10^{-2}$ |
| | 5 | | | | | $2.650 \times 10^{-3}$ |
| | 5 | | GO:BP | GO:0002250 | adaptive immune response | $2.938 \times 10^{-3}$ |
| | 10 | | GO:BP | GO:0002428 | antigen processing and presentation of peptide antigen via MHC class Ib | $3.005 \times 10^{-3}$ |
| | 5 | | | | | $2.561 \times 10^{-2}$ |
| | 10 | | GO:BP | GO:0002474 | antigen processing and presentation of peptide antigen via MHC class I | $3.170 \times 10^{-3}$ |
| | 10 | | GO:BP | GO:0002824 | positive regulation of adaptive immune response based on somatic recombination of immune receptors built from immunoglobulin superfamily domains | $1.200 \times 10^{-2}$ |
| | 5 | | | | | $3.214 \times 10^{-3}$ |
| | 10 | | GO:BP | GO:0002478 | antigen processing and presentation of exogenous peptide antigen | $4.183 \times 10^{-3}$ |
| | 5 | | | | | $9.083 \times 10^{-3}$ |
| | 10 | | GO:BP | GO:0002821 | positive regulation of adaptive immune response | $1.641 \times 10^{-2}$ |
| | 5 | | | | | $4.244 \times 10^{-3}$ |

| | | | | | | |
|---|---|---|---|---|---|---|
| | 10 | | GO:BP | GO:0002819 | regulation of adaptive immune response | $4.032 \times 10^{-2}$ |
| | 5 | | | | | $4.407 \times 10^{-3}$ |
| | 5 | | GO:BP | GO:0002495 | antigen processing and presentation of peptide antigen via MHC class II | $4.462 \times 10^{-3}$ |
| | 5 | | GO:BP | GO:0002504 | antigen processing and presentation of peptide or polysaccharide antigen via MHC class II | $7.425 \times 10^{-3}$ |
| | 10 | | GO:BP | GO:0051251 | positive regulation of lymphocyte activation | $3.070 \times 10^{-2}$ |
| | 5 | | | | | $9.227 \times 10^{-3}$ |
| | 10 | | GO:BP | GO:0019885 | antigen processing and presentation of endogenous peptide antigen via MHC class I | $1.176 \times 10^{-2}$ |
| | 10 | | GO:BP | GO:0019884 | antigen processing and presentation of exogenous antigen | $1.226 \times 10^{-2}$ |
| | 5 | | | | | $2.141 \times 10^{-2}$ |
| | 10 | | GO:BP | GO:0001914 | regulation of T cell mediated cytotoxicity | $1.226 \times 10^{-2}$ |
| | 5 | | | | | $2.141 \times 10^{-2}$ |
| | 5 | | GO:BP | GO:0002399 | MHC class II protein complex assembly | $1.317 \times 10^{-2}$ |
| | 5 | | GO:BP | GO:0002503 | peptide antigen assembly with MHC class II protein complex | $1.317 \times 10^{-2}$ |
| | 10 | | GO:BP | GO:0002475 | antigen processing and presentation via MHC class Ib | $1.417 \times 10^{-2}$ |
| | 10 | | GO:BP | GO:0050870 | positive regulation of T cell activation | $2.291 \times 10^{-2}$ |
| | 5 | | | | | $1.593 \times 10^{-2}$ |
| | 5 | | GO:BP | GO:0050778 | positive regulation of immune response | $1.642 \times 10^{-2}$ |
| | 5 | | GO:BP | GO:0002486 | antigen processing | $1.672 \times 10^{-2}$ |

| | | | | | | |
|---|---|---|---|---|---|---|
| | | | | | and presentation of endogenous peptide antigen via MHC class I via ER pathway, TAP-independent | |
| | 10 | | GO:BP | GO:0002483 | antigen processing and presentation of endogenous peptide antigen | $1.693 \times 10^{-2}$ |
| | 5 | | GO:BP | GO:0002696 | positive regulation of leukocyte activation | $1.799 \times 10^{-2}$ |
| | 5 | | GO:BP | GO:0050776 | regulation of immune response | $1.967 \times 10^{-2}$ |
| | 5 | | GO:BP | GO:0002684 | positive regulation of immune system process | $2.148 \times 10^{-2}$ |
| | 5 | | GO:BP | GO:0050867 | positive regulation of cell activation | $2.524 \times 10^{-2}$ |
| | 10 | | GO:BP | GO:0001913 | T cell mediated cytotoxicity | $2.681 \times 10^{-2}$ |
| | 5 | | | | | $4.022 \times 10^{-2}$ |
| | 10 | | GO:BP | GO:0022409 | positive regulation of cell-cell adhesion | $2.686 \times 10^{-2}$ |
| | 5 | | GO:BP | GO:0002460 | adaptive immune response based on somatic recombination of immune receptors built from immunoglobulin superfamily domains | $2.727 \times 10^{-2}$ |
| | 5 | | GO:BP | GO:0051249 | regulation of lymphocyte activation | $2.757 \times 10^{-2}$ |
| | 10 | | GO:BP | GO:1903039 | positive regulation of leukocyte cell-cell adhesion | $4.616 \times 10^{-2}$ |
| | 5 | | | | | $2.831 \times 10^{-2}$ |
| | 10 | | GO:BP | GO:0050890 | cognition | $3.070 \times 10^{-2}$ |
| | 10 | | GO:BP | GO:1902105 | regulation of leukocyte differentiation | $3.153 \times 10^{-2}$ |
| | 10 | | GO:BP | GO:0051240 | positive regulation of multicellular organismal process | $3.179 \times 10^{-2}$ |
| | 5 | | GO:BP | GO:0002501 | peptide antigen assembly with MHC protein complex | $3.717 \times 10^{-2}$ |

| | | | | | | |
|---|---|---|---|---|---|---|
| | 5 | | GO:BP | GO:0002396 | MHC protein complex assembly | $3.717 \times 10^{-2}$ |
| | 10 | | GO:BP | GO:0048583 | regulation of response to stimulus | $4.235 \times 10^{-2}$ |
| | 10 | | GO:BP | GO:0019883 | antigen processing and presentation of endogenous antigen | $4.860 \times 10^{-2}$ |
| | 10 | | GO:CC | GO:0042611 | MHC protein complex | $8.730 \times 10^{-5}$ |
| | 5 | | | | | $2.358 \times 10^{-6}$ |
| | 10 | | GO:CC | GO:0042613 | MHC class II protein complex | $4.196 \times 10^{-2}$ |
| | 5 | | | | | $4.638 \times 10^{-3}$ |
| | 5 | | GO:CC | GO:0071944 | cell periphery | $8.410 \times 10^{-3}$ |
| | 10 | | GO:CC | GO:0005886 | plasma membrane | $2.881 \times 10^{-2}$ |
| | 5 | | | | | $1.115 \times 10^{-2}$ |
| | 5 | | GO:CC | GO:0098553 | lumenal side of endoplasmic reticulum membrane | $1.542 \times 10^{-2}$ |
| | 5 | | GO:CC | GO:0098576 | lumenal side of membrane | $4.674 \times 10^{-2}$ |
